# A congenital pain insensitivity mutation in the nerve growth factor gene uncouples nociception from affective pain in heterozygous humans and mice

**DOI:** 10.1101/331587

**Authors:** Giovanna Testa, Irene Perini, Marco Mainardi, Chiara Morelli, Francesco Olimpico, Laura Pancrazi, Carla Petrella, Cinzia Severini, Rita Florio, Francesca Malerba, Paul Heppenstall, Mario Costa, India Morrison, Simona Capsoni, Antonino Cattaneo

**Affiliations:** Bio@SNS Laboratory, Scuola Normale Superiore, Pisa, Italy; Department of Clinical and Experimental Medicine, Linköping University, Linköping, Sweden; EMBL Rome, Monterotondo, Italy; Institute of Neuroscience, CNR, Pisa, Italy; Institute of Cell Biology and Neurobiology, CNR, Rome, Italy; Institute of Human Physiology, Department of Medical and Surgical Specialties Sciences, Ferrara, Italy; Neurotrophins and Neurodegenerative Diseases Laboratory, Rita Levi-Montalcini European Brain Research Institute, Rome, Italy

## Abstract

Pain is an unpleasant but necessary sensory experience, which facilitates adaptive behaviours, such as fear. Despite recent advances, the question of how the pain experience influences learning of the fear response is still debated^1,2^. Genetic disorders rendering patients congenitally unable to feel pain have been described, and are usually explained by defects in peripheral nociceptors^3-5^. It is not known how growing up without pain affects central emotional and motivational responses to aversive stimuli. The rare autosomal recessive Hereditary Sensory and Autonomic Neuropathy type V (HSAN V) is caused by a mutation in the nerve growth factor (NGF) gene (R100W). HSAN V homozygous patients display a congenital indifference to painful events, with deficits in peripheral nociceptors but without overt cognitive impairment^6^. In contrast, heterozygous carriers do not present with pain-related deficits and have been identified only through pedigree and genetic screening^7,8^. We exploited this clinically silent population to dissociate nociceptive from affective features of pain. To address this we investigated both heterozygous knock-in mice bearing the R100W mutation (NGF^R100W/m^ mice) and a cohort of HSAN V heterozygous human carriers. Surprisingly, we found that NGF^R100W/m^ mice, despite normal responses to a noxious conditioning stimulus, show a deficit in learned fear while their social and innate fear responses are normal. The lack of pain-related fear response was linked to a reduced activation of anterior cingulate and motor cortices and of striatum, but not in primary somatosensory cortex. Likewise, human heterozygous R100W carriers, despite perceiving noxious stimuli and reporting subjective pain thresholds, show increased reaction latencies in response to painful stimulation, alongside a decreased subjective urgency to react. Functional magnetic resonance imaging (fMRI) revealed, comparably to the mouse data, an altered processing of painful stimuli in rostral anterior cingulate, medial premotor cortical regions and striatum. These findings from both human and mouse HSAN V carriers uncover behavioural and motivational consequences of a mild genetic pain insensitivity on the establishment of pain-dependent affective responses and memories.

---

To shed light on the mechanisms responsible for altered pain perception processing in HSAN V, we generated two mouse knock-in lines harboring (i) the wild type human *ngfb* coding sequence (NGF^h/m^ mice) and (ii) the HSAN V human NGF^R100W^ mutation (Suppl. Fig. 1). Homozygous NGF^R100W/R100W^ mice die by P30 (Suppl. Fig. 2), most likely due to developmental defects caused by the inefficient secretion of the NGF^R100W^ ^5^ (Suppl. Fig. 6a). Indeed, NGF delivery to the pregnant mother and homozygous pups reversed the lethal phenotype of NGF^R100W/R100W^ mice (Suppl. Fig 2). Heterozygous NGF^R100W/m^ mice developed normally and showed an overall mild impairment in nociception, with a reduced sensitivity to capsaicin (Fig. 1a). An increased latency and reduced sensitivity to hot and cold noxious stimuli was observed (Fig. 1b-c), with no change in the pain threshold to hot stimuli, and a decreased temperature sensitivity only above 51°C (Suppl. Fig. 3). NGF^R100W/m^ mice displayed a normal number of reaches in the tape test response (Suppl. Fig. 4a), but a longer response latency (Fig. 1d), correlating with a reduced hairy skin innervation at 6 months of age (and not at 2 months) (Suppl. Fig. 4b). Light touch sensation was normal at both ages tested (Fig. 1e). The lower capsaicin sensitivity of NGF^R100W/m^ mice was fully restored by a long-term treatment with mouse NGF (Fig. 1f).

**Figure 1.**
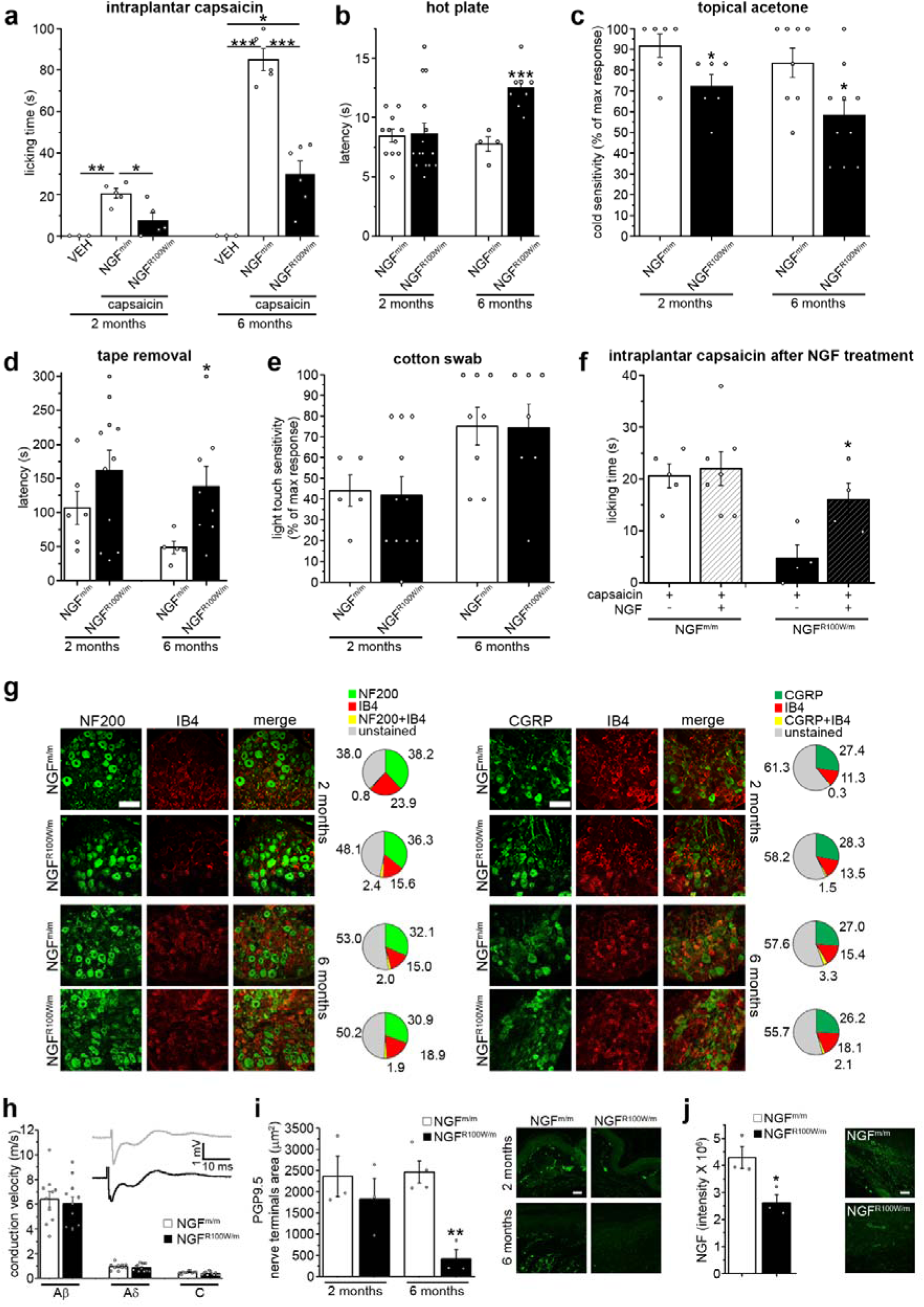
The R100W mutation induces an increased latency to respond to noxious stimuli, normal development of DRG neurons and a reduction of skin innervation. **a,** Decreased hyperalgesic response to capsaicin in juveniles and adults. **b**, Impaired sensitivity to high temperatures in adult HSAN V. **c**, Impaired cold sensitivity in both juveniles and adults. **d**, Decreased hairy skin sensitivity in adult HSAN V mice. **e**, Normal light touch sensitivity. **f**, Rescue of the sensitivity to capsaicin after treatment with mNGF^WT^ from gestation until 2 months of age. **g,** Normal expression of NF200, IB4 and CGRP in DRG neurons in both juveniles and adults (scale bars, 100 µm). **h**, No alteration in conduction velocity of Aβ, Aδ and C fibers in adult mice. **i**, Age-dependent reduction in glabrous skin innervation in HSAN V mice (scale bars, 50 µm). **j**, Decreased NGF levels in the glabrous skin of adult HSAN V mice (scale bars, 50 µm). Data are presented as mean ± SEM.

**Figure 2.**
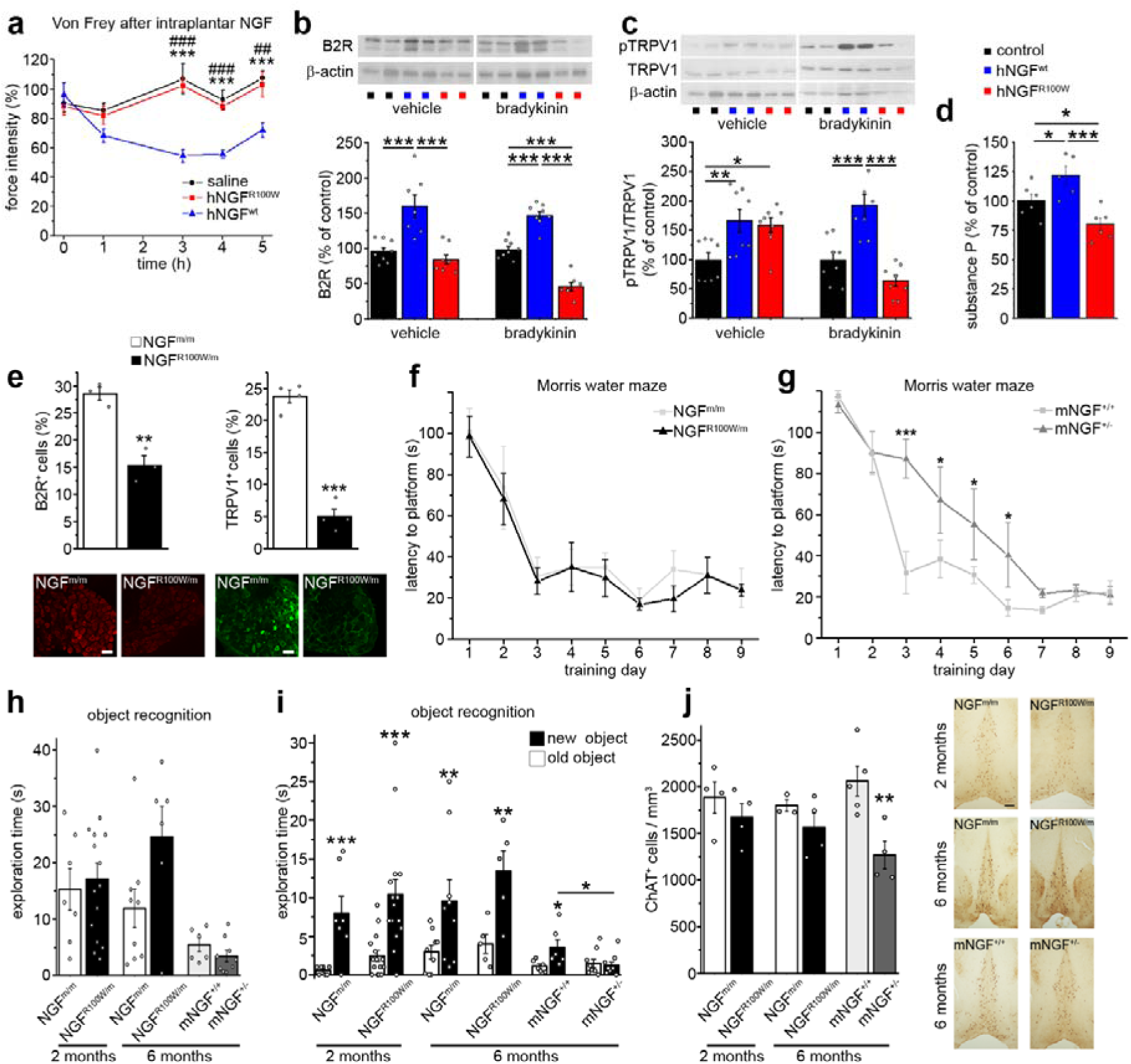
Biochemical and cognitive features of NGF R100W mutation. **a,** Human NGF^R100W^ intraplantar injection causes reduced mechanical sensitization compared to human NGF^WT^. **b**, Downregulation of B2R expression by hNGF^R100W^ in DRG cultures stimulated with bradykinin. **c**, Reduced phosphorylation of TRPV1 by hNGF^R100W^ in DRG cultures stimulated with bradykinin. **d**, Bradykinin-induced SP release in DRG cultures is reduced by hNGF^R100W^ co-treatment compared to hNG^WT^. **e**, Reduced expression of B2R and TRPV1 in DRG neurons of adult mice (scale bars, 50 µm). **f**, Normal Morris water maze learning curve for HSAN V mice. **g**, Delayed Morris water maze learning curve for mNGF^+/-^ mice. **h**, Unimpaired NOR sample phase for HSANV and mNGF^+/-^ mice. **i**, Unimpaired NOR test phase for HSANV and visual recognition memory deficit for mNGF^+/-^ mice. **j**, Normal septal ChAT^+^ neuron density in juvenile and adult HSAN V mice, and decreased density in mNGF^+/-^ mice (scale bars, 200 µm). Data are presented as mean ± SEM.

**Figure 3.**
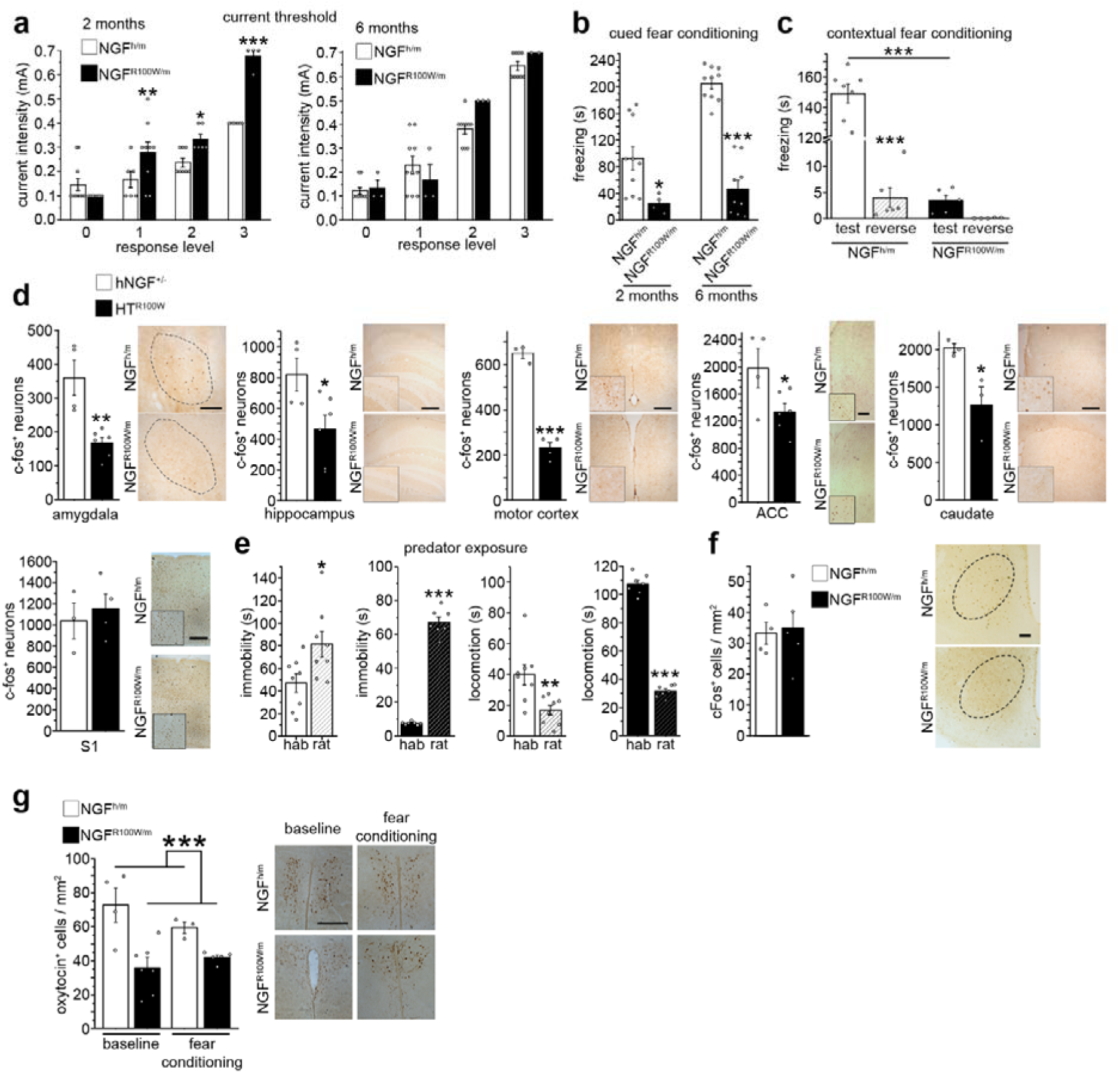
Pain insensitivity impairs acquired but not innate fear. **a**, 2-month-old HSANV mice show higher current intensity threshold for eliciting a maximal response than WT controls, whereas no difference is observed at 6 months of age. **b**, Impaired cued fear memory in HSANV mice. **c**, Reduced freezing in contextual fear conditioning in adult HSAN V mice. **d**, Decreased *c-fos* immunostaining after cued fear conditioning in amygdala, hippocampus, motor cortex, anterior cingulate cortex (ACC), and caudate nucleus of adult HSAN V mice, along with no significant difference in the primary somatosensory cortex (S1) (scale bars, 200 µm, except for ACC, 100 µm). **e**, The hNGF^R100W^ mutation does not affect innate fear and predator exposure reduces locomotion and increases immobility in both mutants and WTs. **f**, No difference in the *c-fos* immunostaining in the hypothalamic VMH of adult WT and HSAN V mice after predator exposure (scale bars, 200 µm). **g**, Reduced oxytocin immunoreactivity in the hypothalamic PVA of NGF^R100W/m^ mice (scale bars, 200 µm).

**Figure 4.**
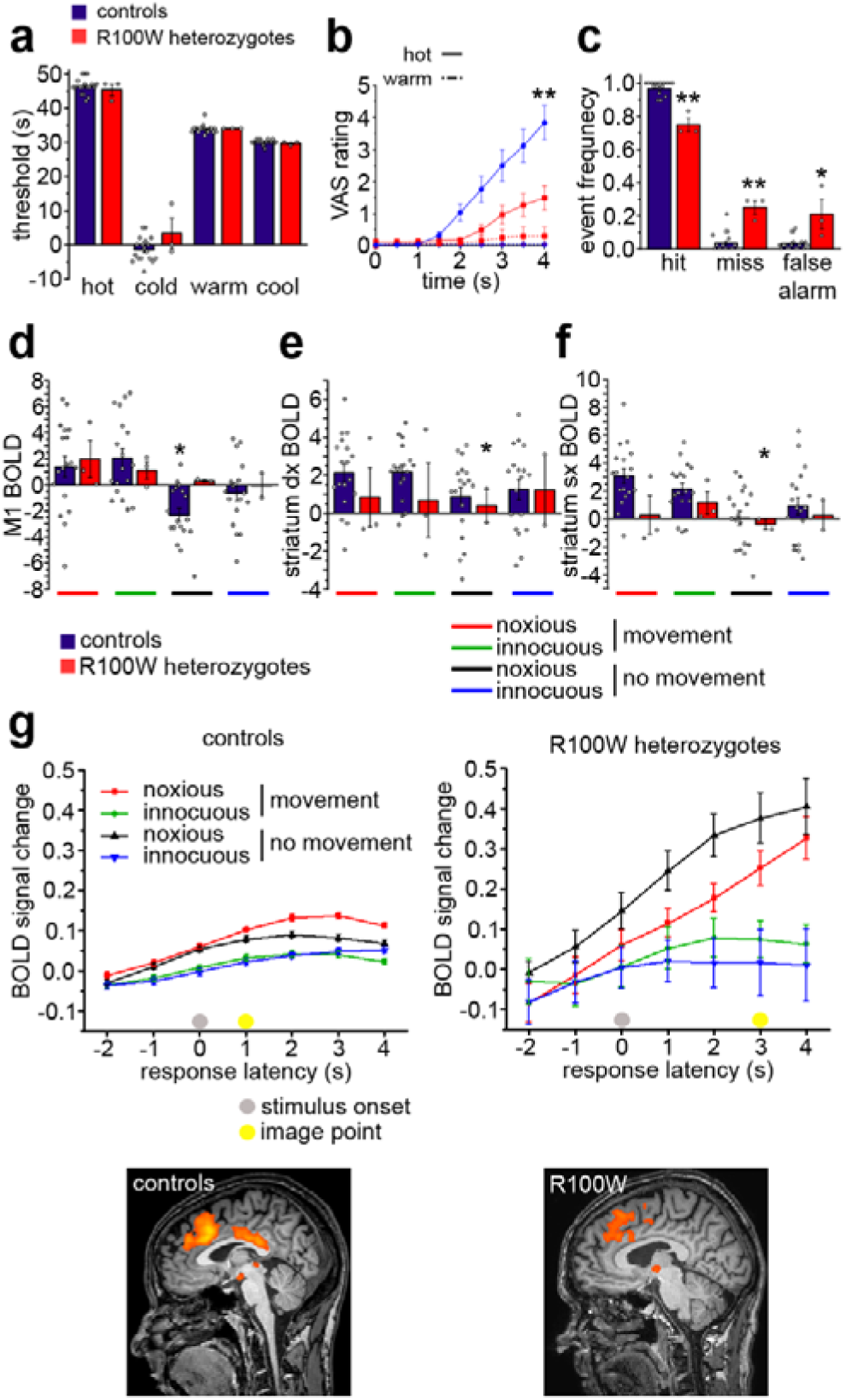
Disrupted modulation and longer response latencies in heterozygous R100W patients. **a,** No differences in discriminative and thermal thresholds between R100W heterozygotes and age-matched controls. **b**, Decreased urge to move in response to painful stimulation in R100W heterozygotes. **c**, Impaired estimation of painful situations in R100W heterozygotes. **d-f**, BOLD responses in motor regions during painful stimulation: no significant modulation in M1 (**d**), and bilateral striatum (**e**, **f**) in R100W heterozygotes; in contrast, control patients show a significant main effect of “movement” vs “no-movement” conditions in M1 and bilateral striatum (**d-f**). **g**, event-related time-course plots of R100W heterozygotes pain-specific BOLD responses in rACC/preSMA show higher latency and loss of baseline return compared to controls. All maps thresholded at p < 0.001 uncorr.

To investigate possible developmental effects on NGF-dependent sensory neurons, we analyzed dorsal root ganglia (DRG) in 2 and 6 months old NGF^R100W/m^ mice. No change was found in the percentage of NF200-, IB4-and of calcitonin gene-related peptide (CGRP)-immunoreactive neurons (Fig. 1g), in line with the unchanged neurotrophic properties of human NGF^R100W^ (hNGF^R100W^) on DRG neurons *in vitro* (Suppl. Fig. 5). Electrical conduction velocities of Aß, Aδ and C sensory fibers were also normal (Fig. 1h). However, glabrous skin innervation decreased with age (Fig. 1i), in correlation with reduced NGF levels in the skin (Fig. 1j).

Next, we explored the ability of hNGF^R100W^ protein to activate and sensitize pain pathways. Mechanical allodynia induced by acute injection of hNGF^R100W^ in the hindpaw of CD1 mice was significantly lower than that induced by wild type hNGF (Fig. 2a). Incubation of DRG neurons with hNGF^R100W^, followed by acute administration of the inflammatory neuropeptide bradikinin (BK)^9^, leads to reduced bradykinin B2R receptor expression (Fig. 2b), TRPV1 phosphorylation (Fig. 2c) and Substance P release (Fig. 2d). Thus, priming of DRG neurons with hNGF^R100W^ diminished their propensity to sensitize. Consistently, expression of B2R and TRPV1 was reduced in DRG from 6-month-old NGF^R100W/m^ mice (Fig. 2e). Thus, hNGF^R100W^ protein displays an indistinguishable neurotrophic potency from wild type hNGF, while showing a reduced ability to sensitize sensory neurons. This extends data showing that hNGF^R100W^ uncouples trophic effects from nociceptive functions of NGF^10-12^.

This prompted us to investigate whether the hNGF^R100W^ mutation might affect learning and memory behaviours triggered by painful events. To control for general learning and memory deficits in NGF^R100W/m^ mice, that might result from the impaired secretion of hNGF^R100W^ and of wild type NGF, when coexpressed with hNGF^R100W^ (Suppl. Fig. 6), we performed Morris Water maze, novel object recognition and Y maze tests, in comparison to heterozygous murine NGF mice (mNGF^+/-^)^13^. NGF^R100W/m^ mice showed no learning and memory deficits in any of these tests and a normal number of Choline Acetyl-Transferase (ChAT)-immunoreactive NGF target neurons (Fig. 2f-j and Suppl. Fig. 7). On the contrary, mNGF^*+/-*^ mice showed, as expected^14^, a deficit in the Morris Water maze (Fig. 2g) and in the object recognition tests (Fig. 2h,i), together with a reduction of ChAT-immunoreactive neurons in the medial septum of the basal forebrain (Fig. 2j). Thus, the hNGF^R100W^ mutation, unlike *ngf* gene deletion, does not affect spatial learning and memory processes, in keeping with heterozygous R100W human carriers, who are reported to be cognitively normal^6^. In contrast, NGF^R100W/m^ mice showed an age-related reduction of anxiety, as revealed by alterations in the elevated plus maze, marble burying and nest building tests (Suppl. Fig. 8). We reasoned that this might reflect NGF^R100W/m^ mice being less anxious about unexpected potentially harmful events. We therefore asked whether central processing of pain-related events and memory processes might be affected in these mice, despite their mild peripheral pain phenotype.

To this purpose, we used the fear conditioning test^15^. The intensity of the foot shock stimulus was adjusted to reach the maximum vocalization response in NGF^R100W/m^ and NGF^h/m^ mice (Fig. 3a). The behavioural response to the shock was identical in the two groups. Surprisingly, NGF^R100W/m^ mice, unlike control NGF^h/m^ mice, did not show a fear freezing response when presented with the auditory cue alone (recall phase of the test), despite normal freezing responses in the association phase (Fig. 3b). The reduced freezing response was not due to an auditory deficit, since NGF^R100W/m^ mice showed a reduced freezing response also in the contextual fear conditioning test (Fig. 3c). We conclude that NGF^R100W/m^ mice are not able to associate the conditioning stimulus to the noxious unconditioned stimulus and do not interpret the former as an event potentially leading to pain, despite a normal response to the shock per se.

We compared which brain areas are activated after exposure to the cue, in wild type versus NGF^R100W/m^ mice. The number of c-fos-immunoreactive neurons activated in NGF^R100W/m^ mice during the recall phase was reduced in the amygdala, hippocampus, motor cortex, anterior cingulate cortex and anterior caudate nucleus, with respect to NGF^h/m^ mice, but, importantly, not in the primary somatosensory cortex (Fig. 3d).

Is the observed lack of fear response in NGF^R100W/m^ mice specific for painful stimuli? When NGF^R100W/m^ mice were exposed to a predator (Fig. 3e) or to a stranger mouse (Suppl. Fig. 9), a paradigm of innate fear not related to pain, they behaved identically to wild type mice. Consistently, after exposure to the predator, the c-fos activation pattern in the ventromedial hypothalamic nucleus, which mediates this specific fear response^16^, is identical to control NGF^h/m^ mice (Fig. 3f). We conclude that NGF^R100W/m^ mice have a deficit specific to learned responses triggered by painful events, despite only having a mildly impaired peripheral nociception. The results suggest that the reduced behavioural response to the painful stimulus is caused by a specifically altered interpretation of the pain-relevant signal.

In search for a molecular mediator of this reduced fear response we focused on oxytocin (OXT), a molecule known to influence fear memory formation^17^. In NGF^R100W/m^ mice, plasma OXT was reduced (Suppl. Fig. 10). Moreover, the number of OXT-immunoreactive neurons in the paraventricular hypothalamic nucleus was greatly decreased, before and after fear conditioning (Fig. 3g). This finding is consistent with observations in OXTR^FB/FB^ mice, which show a reduced freezing response after cued fear conditioning^18^ and suggests that reduced OXT signaling mediates the impaired fear response in NGF^R100W/m^ mice.

To relate the mouse findings to the human heterozygous R100W phenotype, discriminative and pain thresholds for heat and cold were investigated in heterozygous R100W carriers and normal controls. Heterozygotes have variable alterations in the density of C-fiber innervation, which overlap with the general population^8^. No significant differences in mean pain threshold values were found between the two groups (Fig. 4a). The subjective motivational responses to above-threshold painful thermal stimuli were measured by an “urge to move” rating^19^. The R100W carrier group showed a significantly lower urge to move, with higher response latencies (Fig. 4b) and a significantly higher error frequency in a pain/no pain discrimination task (Fig. 4c). Using fMRI, carriers and healthy controls underwent painful vs innocuous thermal stimulation during scanning. The carrier group showed a greatly reduced blood-oxygen-level-dependent (BOLD) activation in primary motor cortex (M1) and striatum, compared to controls (Fig. 4d-f), but a normal activation in primary somatosensory cortex (data not shown). During the first seconds following the onset of the thermal stimulus, controls, but not carriers, showed a main effect of pain stimulus in rostral anterior cingulate cortex (rACC) and in pre-supplementary motor area (pre-SMA). Time course analysis of the full 4-second trial revealed that, although this main effect did not manifest at stimulus onset in R100W carriers, these regions did show a delayed activation by the last second of the trial (Fig. 4g). Unlike rACC/SMA responses in controls, this activation in carriers did not return to baseline at the end of the trial (Fig. 4g). Altogether, these human data reveal a comparable impairment to NGF^R100W/m^ mice in integrating pain-related afferent signalling with task-dependent behavioural responses, and a dysregulation of pain-triggered motor-related responses, despite normal pain thresholds and activation of primary somatosensory cortex.

Painful stimuli have a number of different attributes, including peripherally encoded sensory-discriminative (such as location, intensity and quality), as well as centrally encoded affective and emotional valence (such as unpleasantness)^20^. The mild nociceptive impairment observed in HSAN V heterozygous mice and human carriers allowed us to demonstrate that no major developmental defect occurs, even if the mutant R100W protein interferes with the secretion of the wild type protein (possibly by forming NGF R100W/WT heterodimers; Suppl. Fig. 6a). This gave us the opportunity to study whether and how the peripheral sensory input of pain signals can be dissociated from the central elaboration of the affective features of the pain stimulus, in both humans and mice. We found that human and mice HSAN V heterozygous carriers display alterations in central behavioural pain responses, showing that even with normal afferent nociception, the meaning and interpretation of a painful stimulus as an unpleasant feeling to avoid, can be severely affected, to the extent of changing behavioural affective responses. Thus, the central elaboration of pain can be dissociated from the peripheral sensory nociception.

We identified a common set of mouse and human brain regions, that are differentially activated in heterozygous HSAN V vs the corresponding controls: the ACC, SMA - medial cortical regions and striatal areas, involved in premotor preparation of motivated behaviour. Activity in the ACC has been previously suggested to be related to the affective features of pain, such as subjective feelings of unpleasantness^21^, and is tightly linked to producing behavioural responses to pain^19^.

Painful stimuli induce robust learning and memory formation^22-24^. The mouse data reveal a selective defect in the ability to form a pain-related fear memory, indicating an inability to form episodic memories triggered by painful events. In contrast, predator and social fears are not affected. The latter forms of fear, even though they also represent pain-relevant situations, are not learned, but innate^25^. The observed age-related alterations of anxiety-like behaviours in NGF^R100W/m^ mice are consistent with a lack of pain-related fear learning. This leads us to conclude that the NGF^R100W^ mutation specifically affects the expression of learned, pain-related behaviours, without affecting other forms of learning. A candidate molecular mediator of the reduced fear response in NGF^R100W/m^ mice, and possibly for the reduced urge to move response in humans, is oxytocin, a well-known bidirectional modulator of both fear and pain related memories^17^. It remains to be determined if hNGF^R100W^ modulates OXT expression, directly or indirectly.

Congenital insensitivity to pain is generally explained by a peripheral block of nociception^4^. For instance, the painlessness phenotype deriving from a Na_v_1.7 mutation could be reversed by the administration of the opioid antagonist naloxone in patients and mice, by potentiating noxious peripheral input to the spinal cord^26^. We focused, instead, on the central aspects of pain perception, and our results demonstrate that the effects of NGF on pain go beyond its known actions on peripheral nociception^27-29^ and uncover a new action at the brain level.

Taken together, the human and mouse results converge to demonstrate the disruption by the hNGF^R100W^ mutation of a centrally-mediated system that integrates sensory noxious information with sensory cues to generate an adaptive response, and reveal a defect in interpreting painful stimuli as relevant for determining adaptive behavioural responses. By studying the effects of the hNGF^R100W^ mutation in the heterozygous state, we have opened a window on the subjective experience of unpleasantness related to pain stimulation, linking it to a precise brain system and candidate molecules, an approach that could be relevant to design much needed strategies to treat chronic pain states.

## Acknowledgements

The authors would like to thank Lorenza Ronfani, Rosanna Rinaldi, Ivana Benzoni (San Raffaele Hospital s.r.l.), Maria Antonietta Calvello, Vania Liverani, Nicola Maria Carucci, Francesco Gobbo, Caterina Rizzi, Alessandro Viegi and Alexia Tiberi (all at Scuola Normale Bio@SNS lab) for their help and support, and Andrea Poli (Institute of Neuroscience, CNR) for the initial instructions on fear conditioning test. This work was supported by the EU FP7 PAINCAGE project (to AC, grant agreement n. 603191) and by Telethon (to AC, grant n. GGP11179).

## Author Contributions

Generation of genetically engineered mouse model: G.T, L.P, and M.C Behavioural studies: G.T, M.M and F.O with support from S.C; Human data: I.P and I.M; Histochemical and biochemical analysis in mice and cells: G.T, F.O M.M. with support from S.C. DRG, skin and conduction velocity analysis: C.M, G.T with support from P.H. In vitro studies: C.P and C.S. Production and purification of hNGF^R100W^ protein: F.M and R.F. G.T analysed the results, and contributed to discussions and manuscript preparation. S.C and A.C directed the project, supervised data analysis, and wrote the manuscript.

